# Centrin-deficient *Leishmania mexicana* confers protection against Old World visceral leishmaniasis

**DOI:** 10.1101/2022.06.27.497754

**Authors:** Subir Karmakar, Greta Volpedo, Wen-Wei Zhang, Patrick Lypaczewski, Nevien Ismail, Fabiano Oliveira, James Oristian, Claudio Meneses, Sreenivas Gannavaram, Shaden Kamhawi, Shinjiro Hamano, Jesus G. Valenzuela, Greg Matlashewski, Abhay R Satoskar, Ranadhir Dey, Hira L. Nakhasi

## Abstract

Leishmaniasis is one of the top neglected tropical diseases with significant morbidity and mortality in low and middle-income countries (LMIC). However, this disease is also spreading in the developed world. Currently, there is a lack of effective strategies to control this disease. Vaccination can be an effective measure to control leishmaniasis and has the potential to achieve disease elimination. Recently, we have generated *centrin* gene-deleted new world *L. mexicana* (*LmexCen*^*-/-*^) parasites using CRISPR/Cas9 and showed that they protect mice against a homologous *L. mexicana* infection that causes cutaneous disease. In this study, we tested whether *LmexCen*^*-/-*^ parasites can also protect against visceral leishmaniasis caused by *L. donovani* in a hamster model. We show that immunization with *LmexCen*^*-/-*^ parasites is safe and does not cause lesions. Furthermore, such immunization conferred protection against visceral leishmaniasis caused by a needle-initiated *L. donovani* challenge, as indicated by a significant reduction in the parasite burdens in the spleen and liver and lack of mortality. Similar control of parasite burden was also observed against a sand fly mediated *L. donovani* challenge. Importantly, immunization with *LmexCen*^*-/-*^ down-regulated the Th2 response as indicated by a significant reduction in the anti-inflammatory cytokines such as IL-10 and IL-4 and increased pro-inflammatory cytokine IFN-γ resulting in higher IFN-γ/IL-10 and IFN-γ/IL4 ratios compared to non-immunized animals. This contrasts with our studies with *L. major centrin* deletion mutants that showed a dominant Th1 response compared to *L. major* wild-type infection suggesting the divergent mechanisms of protection in the two mutant parasites. *LmexCen*^*-/-*^ immunization resulted in long-lasting protection against *L. donovani* infection. Further, since the efficacy of *LmCen*^*-/-*^ has not been determined against *Leishmania* strains prevalent in the Americas, *LmexCen*^*-/-*^ may be a viable alternative. Taken together, our study demonstrates that immunization with *LmexCen*^*-/-*^ parasites is safe and efficacious against old world visceral leishmaniasis.

## INTRODUCTION

Leishmaniases are vector-borne parasitic diseases affecting millions of people globally, which clinically range from self-healing cutaneous (CL) to systemic and often fatal visceral leishmaniasis (VL) ^1,2^. The various clinical forms of leishmaniases are caused by different parasite species, and CL is caused by the infections with *L. major* or *L. tropica* in the old world and with *L. mexicana* species complex in the new world. The more severe and life-threatening VL is caused by the *L. donovani* complex in the old world and by *L. infantum* (also known as *L. chagasi*) in the new world ^3^.

Since most of the chemotherapies against leishmaniasis suffer from limitations like toxicity, high cost, the necessity for long-term use and most importantly emergence of drug resistance ^4-6^, vaccination would be effective in achieving control and elimination of Leishmaniasis. Because recovery from primary *Leishmania* infection gives lifelong protection from future infections, leading to the insight that vaccination is feasible against leishmaniasis ^7-10^. However, there is no licensed vaccine available for human use against any form of leishmaniasis.

Earlier studies have shown that deliberate infections with a low dose of virulent live wild-type dermotropic *L. major* parasites confers protection against reinfection, a process termed leishmanization ^11-13^. Leishmanization also afforded cross-protection against VL in various animal models and humans ^8,14,15^. However, such method of immunization is not practical because of the safety concerns regarding skin lesions that last for months at the site of inoculation in a naïve population ^16,17^. In contrast, immunization with live attenuated dermotropic *Leishmania* parasites which are non-pathogenic and provide a complete array of antigens like their wild-type analogues, could be a promising vaccine candidate against both CL and VL. While leishmanization with old world species of *L. major* is widely studied to identify the mediators of protective immunity and used as a standard to replicate in numerous experimental vaccine studies, there is no equivalent leishmanization with the new world species of *Leishmania*, such as *L. mexicana*. Infections with *L. mexicana* species cause a more chronic pathology that, unlike skin lesions caused by *L. major* infection, does not self-resolve. Further, infection with *L. mexicana* presents distinct clinical features and pathologies to old world *Leishmania* species. For example, *L. mexicana* infection, which initially causes localized lesions, can progress into diffuse lesions in other parts of the body away from the site of infection, with no delayed-type hypersensitivity (DTH) response ^18,19^. In addition, induction of a Th2 dominant response is responsible for pathogenesis in *L. mexicana* infection of mice ^20^. A similar dominant Th2 response has been shown to play an important role in the pathogenesis of a related new world species, *L. braziliensis*, in a hamster model ^21^. Therefore, it is necessary to develop a vaccine candidate that is effective at preventing infections with new world *Leishmania* species. We recently demonstrated the safety and efficacy of a marker-free *centrin* gene-deleted live attenuated new world dermotropic *L. mexicana* (*LmexCen*^*-/-*^) parasite in a mouse model of new world cutaneous leishmaniasis ^22^. Centrin is a calcium-binding protein, essential in the duplication of centrosomes in eukaryotes including *Leishmania* ^23,24^. *Centrin* gene-deficient *Leishmania* parasites are replication-deficient only in the intracellular amastigote stage but can be easily grown in promastigote culture ^23^. Similar deletion of centrin in *L. major* (*LmCen*^*-/-*^) shows that immunization with this vaccine derived from the old world species of *Leishmania* can protect against CL and VL challenge infections ^25,26^.

The current study examined whether a new world dermotropic *LmexCen*^*-/-*^ parasite is similarly effective against fatal VL in a hamster model. Hamsters develop clinicopathological symptoms of VL similar to human VL, including succumbing to death and are considered a gold standard model of VL ^27^. Data showed that immunization of hamsters with live attenuated *LmexCen*^*-/-*^ is safe and cross-protects against a *Leishmania donovani* challenge infection. Moreover, *LmexCen*^-/-^ vaccinated hamsters showed downregulation of Th2 response as indicated by reduced IL-10 and IL-4 cytokine expression and higher IFN-γ response resulting in higher IFN- γ/IL-10 and IFN-γ/IL-4 ratios, the key biomarkers of protection compared to *LmexWT* infection. Further, immunization with *LmexCen*^*-/-*^ resulted in long-lasting protection against *L. donovani* challenge infection through needle injection or an infected sand fly. These studies show that immunization with genetically modified new world *Leishmania mexicana* parasites has the potential as a vaccine against new world *Leishmania* species but also old world visceral leishmaniasis.

## RESULTS

### *LmexCen*^*-/-*^ parasites do not cause any lesion development in hamsters

To evaluate the non-pathogenicity of *LmexCen*^*-/-*^ parasites as a vaccine candidate, hamsters were injected intradermally with 10^6^ *LmexCen*^*-/-*^ promastigotes. Wild-type *L. mexicana* (10^6^, *LmexWT*) parasites injected-hamsters were used as a control group. Lesion development was monitored up to 11-weeks post-injection, and parasite loads were determined at this study endpoint through the serial dilution method (Fig. 1A). Hamsters injected with *LmexCen*^*-/-*^ parasites did not develop any visible lesions up to 11-weeks of post-injection (Fig. 1B, C). In contrast, hamsters injected with *LmexWT* parasites developed ear lesions within 5-weeks of injection that progressively increased in size (Fig. 1B, C). Low levels of viable parasites were recovered from the ears (2 out of 7) and draining lymph nodes (1 of 7) of hamsters (Fig. 1D, E). The *LmexWT*-injected hamsters had significantly higher parasite loads both in the ears (∼10^6^ Fig. 1D) and in the dLNs (∼10^4^ Fig. 1E) compared to *LmexCen*^*-/-*^ injected animals. No viable parasites were recovered from the spleen, liver, or bone marrow of either *LmexCen*^*-/-*^ or *LmexWT* injected hamsters at 11-weeks post-inoculation (data not shown). These results show that *LmexCen*^*-/-*^ parasites are non-pathogenic, thus safe.

**Figure 1.**
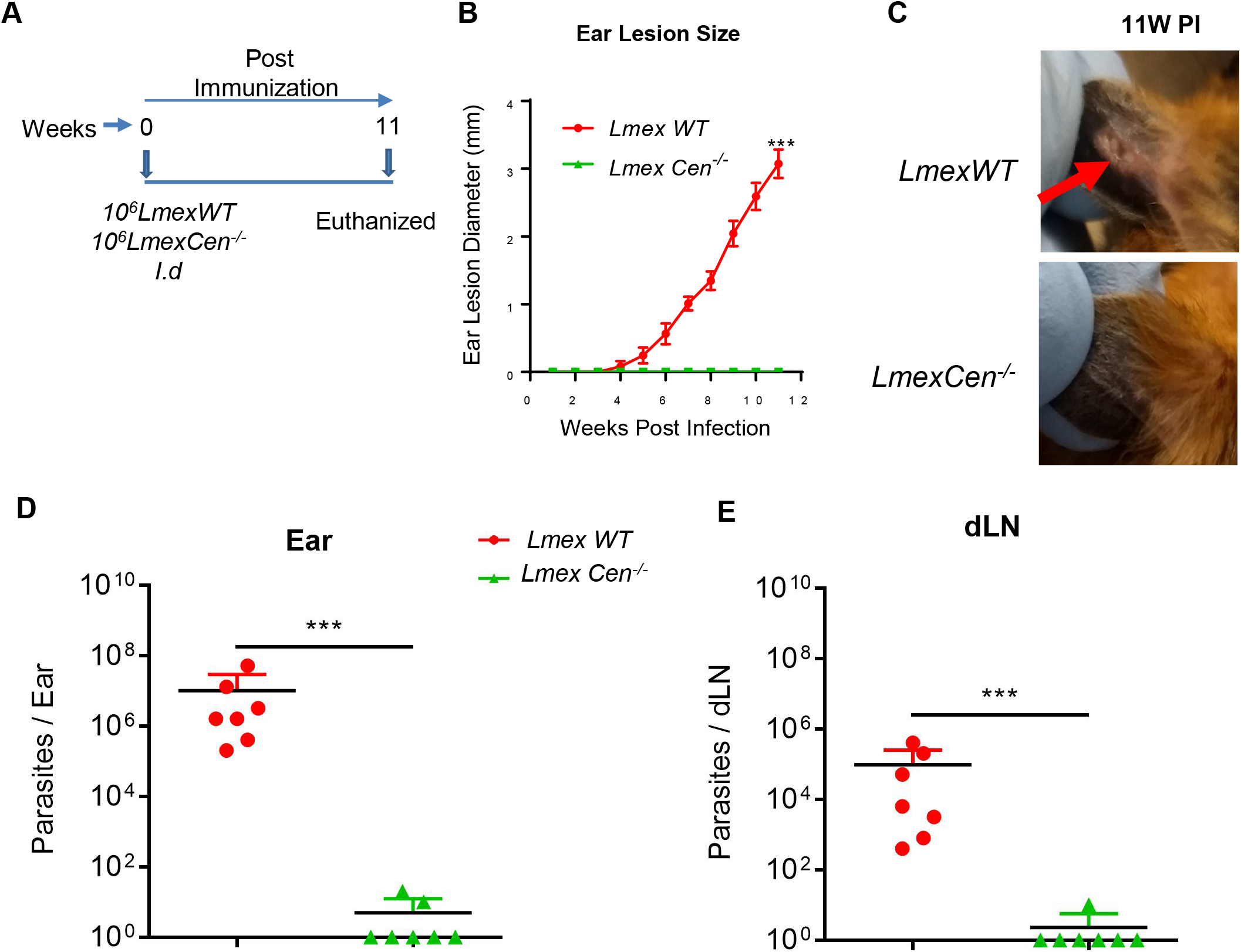
Live attenuated *LmexCen*^*-/-*^ parasites are safe and do not cause lesions in hamster. **(A)** Schematic representation of the experimental plan. (**B**) Lesion size was monitored every week in hamsters injected with 10^6^-total stationary phase either *LmexWT* or *LmexCen*^*-/-*^ parasites by intradermal (ID) injection. Ear lesion diameters were measured at indicated weeks post inoculation. Results (SEM) are representative cumulative effect of two independent experiments, 1 ear, total 7 hamsters per group (***p-0.0006 values were determined by Mann Whitney two-tailed test). (**C**) Photographs of representative ears of *LmexWT* and *LmexCen*^*-/-*^ immunized hamsters at 11 weeks post inoculation (PI). Red arrow indicates the lesion development. (**D, E**) Parasites load in the ear (D) and draining lymph node (dLN) (E) of *LmexWT* and *LmexCen*^*-/-*^ immunized hamsters were determined by serial dilution assay at 11 weeks (n=7/group of hamsters) post inoculation. Results (Mean ± SD) represent cumulative of two independent experiments (***p-0.0006 values were determined by Mann Whitney two-tailed test).

### Immunization with *LmexCen*^*-/-*^ induces a pro-inflammatory immune response

Next, we evaluated the immune response associated with the immunization by analysing the expression of major pro-(IFN-γ) and anti-(IL-10 and IL-4) inflammatory cytokines in the splenocytes of *LmexCen*^*-/-*^ and *LmexWT* injected hamsters at 11 weeks post-infection (Fig.2). To measure antigen specific immune response, splenocytes were either unstimulated or stimulated with *L. mexicana* freeze thawed antigen (±FTAg). In both *LmexCen*^*-/-*^ immunized and *LmexWT* infected animals, upon antigen stimulation, expression of IFN-γ increased in the spleen even though the increase was not statistically significant between the two groups (Fig. 2A). Of note, the parasite burden was significantly different between the two groups (Fig. 1 from parasite burden). We also measured other inflammatory cytokines TNF-α and IL-12p40 at this time point (Fig. 2B-C). Our data indicate no significant difference in their expression between *LmexWT* and *LmexCen*^*-/-*^ immunized groups. Measurement of anti-inflammatory cytokines IL-10 and IL-4, on the other hand, revealed contrasting results. In the spleens of *LmexCen*^*-/-*^ immunized hamsters both the anti-inflammatory cytokines, IL-10 and IL-4, were significantly inhibited compared to *LmexWT* infected group (Fig. 2D-E). Moreover, the ratios of IFN-γ to IL-10 as well as IFN-γ to IL-4 were significantly higher in the spleens of the *LmexCen*^*-/-*^ immunized group compared to *LmexWT* injected animals (Fig. 2F-G), suggesting that *LmexCen*^*-/-*^ immunization induces a pro-inflammatory environment in the spleens of immunized animals. Collectively, these results demonstrate that the live attenuated *LmexCen*^*- /-*^ parasites are immunogenic in a hamster model in the absence of cutaneous lesions.

**Figure 2.**
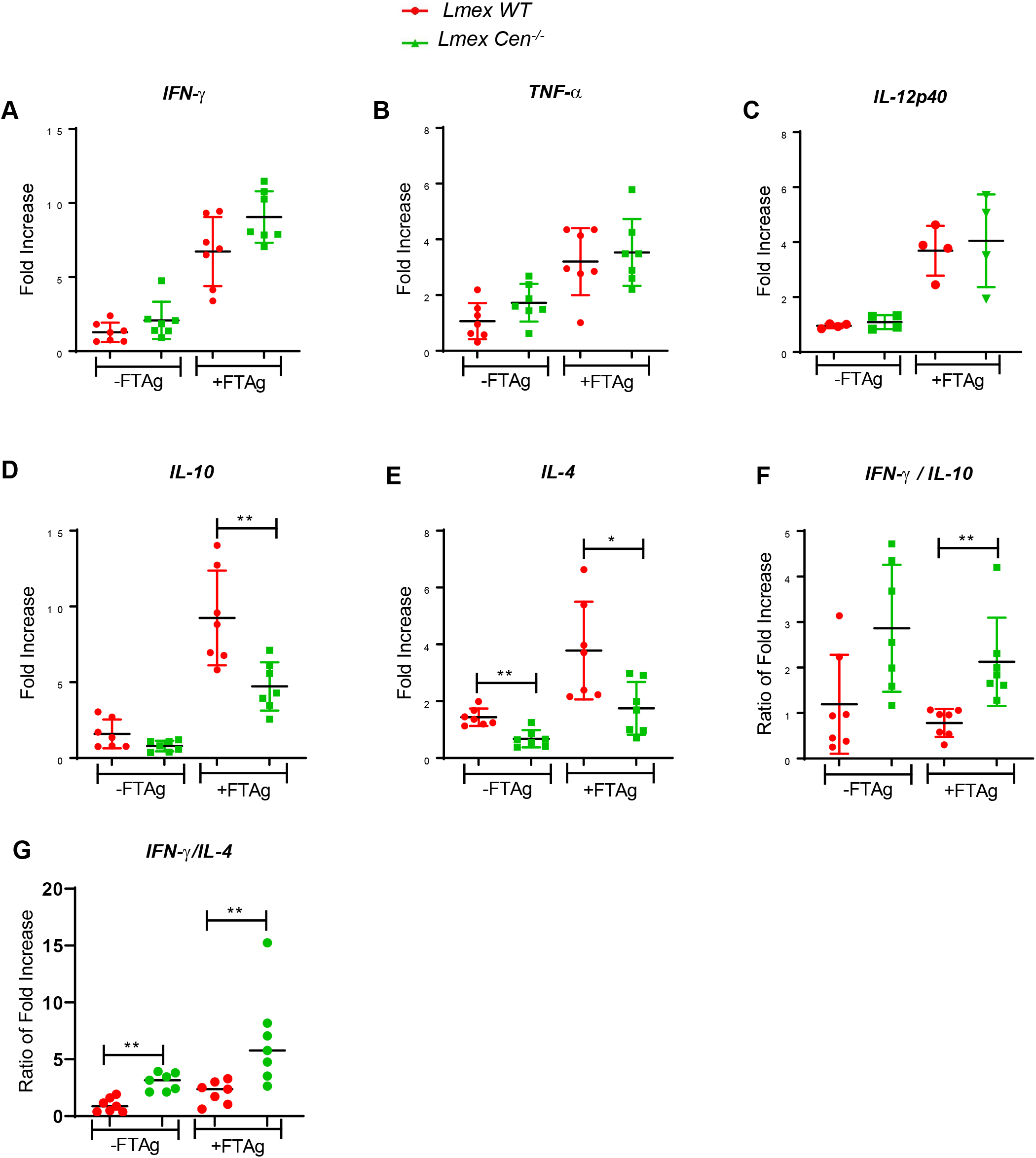
Live attenuated *LmexCen*^*-/-*^ parasites down regulate Th2 cytokine expression in spleen of hamsters. **(A)** IFN-γ, (**B**) TNF-α, (**C**) IL-12p40 (**D**) IL-10, and (**E**) IL-4 expression in the spleen of *LmexWT* or *LmexCen*^*−/−*^ inoculated hamsters (n=7/per group) was evaluated by RT-PCR following 11 weeks post-inoculation. The ratio of IFN-γ/IL-10 (**F**) and IFN-γ/IL-4 (**G**) expression in the spleen was also determined. Except IL-12p40, results (mean ± SD) are cumulative of two independent experiment. Statistical analysis was performed by Mann- Whitney two-tailed test.

### Immunization with *LmexCen*^*-/-*^ protects against challenge infection with *L. donovani* through needle and sand fly mediated infection in a hamster model

Since *LmexCen*^*-/-*^ parasites are immunogenic in hamsters, we investigated the efficacy of *LmexCen*^*-/-*^ immunization against challenge infection with *L. donovani*. Hamsters were intradermally immunized with *LmexCen*^*-/-*^ parasites. Immunized hamsters were challenged at 11-weeks post-immunization with virulent *L. donovani* wild-type (*LdWT*) parasites and monitored up to 15 months post-challenge (Fig. 3A). Survival data revealed that *LmexCen*^*-/-*^ immunization protected 80% of animals against mortality associated with *L. donovani* infection, as demonstrated by the survival of 4- out of the 5- animals that remained healthy up to 15- months post-challenge, the endpoint of our study. All the age-matched non-immunized- hamsters died with symptoms characteristic of human VL starting from 9 months post- challenge timepoint and up to–15 months corresponding to the endpoint of the study following *L. donovani* challenge (Fig. 3B). Spleens isolated from the moribund hamsters from the non-immunized challenged group demonstrated splenomegaly (Fig 3C). In contrast, spleens from *LmexCen*^*-/-*^ immunized challenged group showed no splenomegaly consistent with a lack of pathology in the immunized hamsters. Moreover, analysis of the parasite burden revealed significant control of parasite burden both in the spleen (∼5 log reduction, Fig. 3D) and liver tissues (∼4 log reduction, Fig. 3E) of *LmexCen*^*-/-*^ immunized hamsters compared to the nonimmunized-challenged group. Of note, 60% (3 out of 5) of the livers from immunized animals had no detectable parasites.

**Figure 3.**
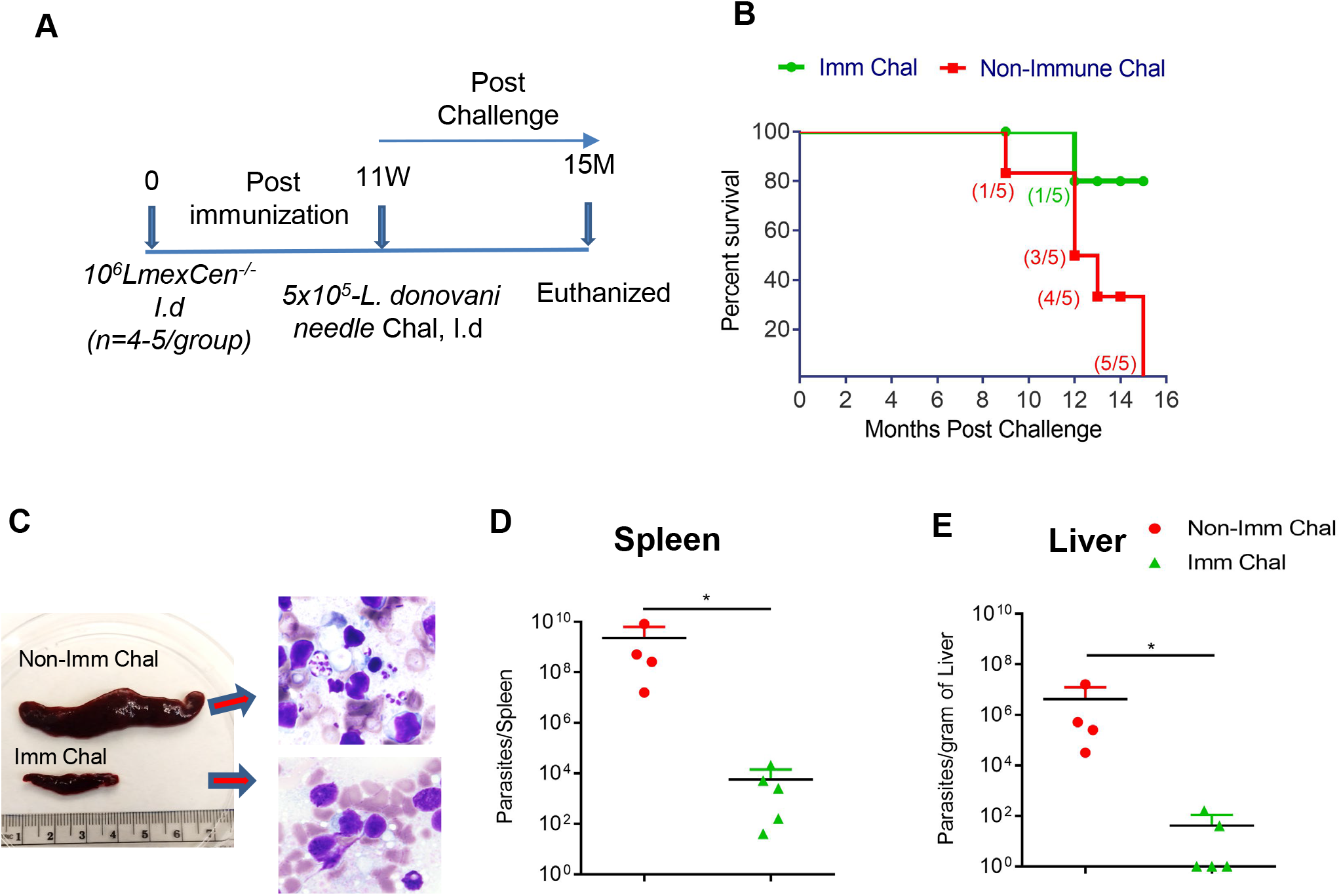
*LmexCen*^*−/−*^ immunization confers protection against *L. donovani* through needle challenge in hamsters. **(A)** Schematic representation of the experimental plan to determine the protection efficacy of *LmexCen*^*-/-*^ parasite. (**B**) Kaplan-Meier survival curves of *LmexCen*^*−/−*^ *-*immunized hamsters (Imm Chal; green lines, n=5) following challenge with *L. donovani* infected sand flies and compared with age-matched non-immunized challenged group (Non Imm Chal; red lines, n=5).). (**C**) Photographs (left panel) of representative one spleen samples of both *LmexCen*^*-/-*^*-* immunized and non-immunized hamsters following 12months of post-needle challenge hamster is shown. (**D** and **E**) Parasite loads in the spleen (D) and liver (E) of hamsters either immunized with *LmexCen*^*-/-*^ (Imm Chal, n=5) parasites or age-matched non-immunized control (Non-Imm Chal, n=4) were determined following 9-15 months of post-needle challenge with *L. donovani*. Results (mean ± SD) are from one experiment. Statistical analysis (*p-0.01) was performed by Mann-Whitney two-tailed test

To confirm if immunization with *LmexCen*^*-/-*^ can similarly protect hamsters against a more rigorous challenge with a sand fly mediated *L. donovani* infection, immunized hamsters were exposed to *L. donovani* infected-sand flies at 11 weeks post-immunization and monitored for 12 months post-challenge (Fig. 4A). The animals were sacrificed at 10-12 months, and parasite burden in spleens and livers was determined. There was a significant reduction of parasite burden in both the spleen and liver of immunized hamsters compared to non-immunized animals (∼4 log reduction Fig. 4B-C). Together, these data demonstrate that *LmexCen*^*-/-*^ immunization protects against both needle and sand fly mediated infection in a hamster model of VL, suggesting that immunization with the new world CL vaccine can also protect old world VL.

**Figure 4.**
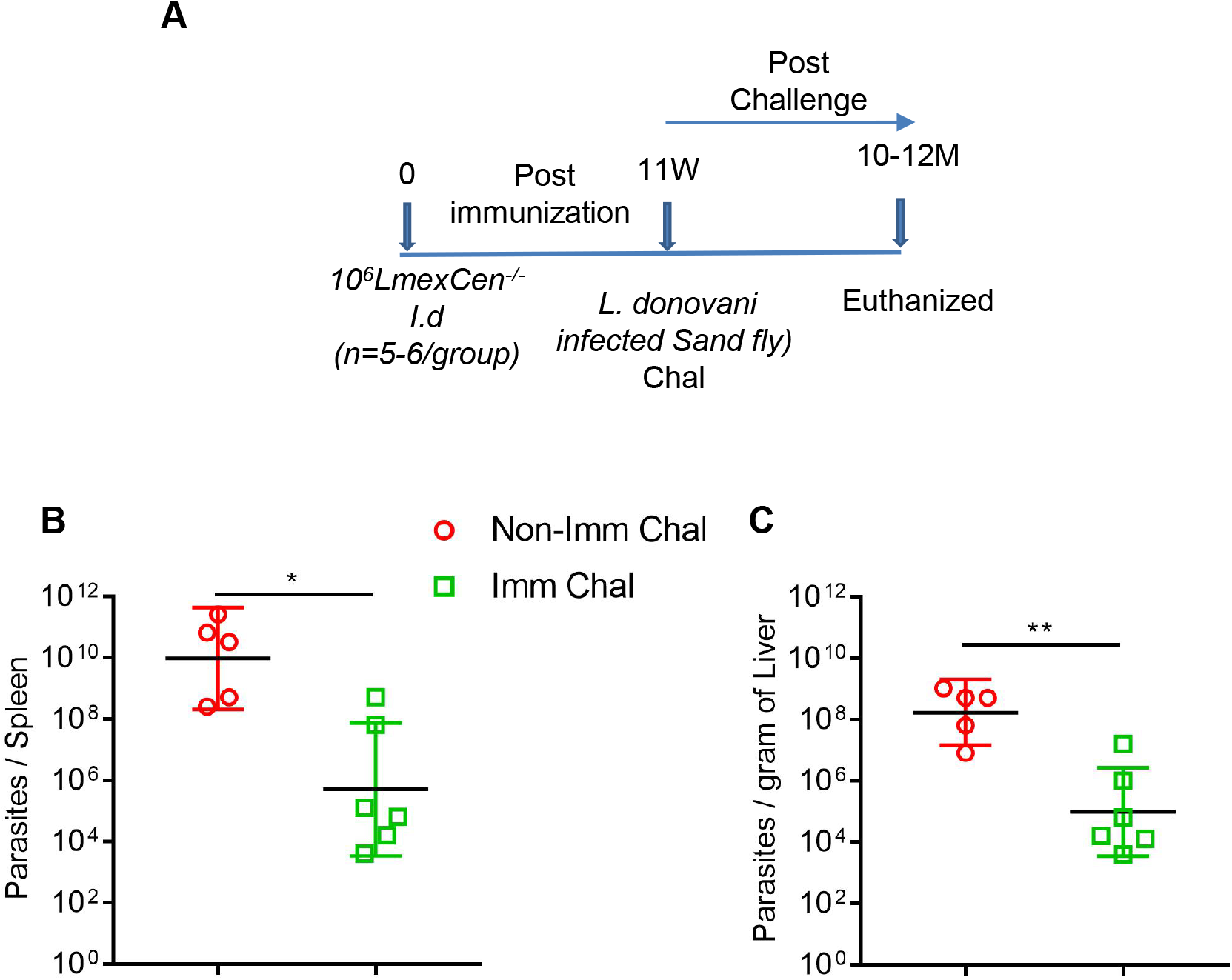
*LmexCen*^*−/−*^ immunization confers protection against sand fly transmitted *L. donovani* infection in hamsters. **(A)** Schematic representation of the experimental plan. Parasite load in the spleen (B) and liver (C) of age matched non-immunized (Non-Imm chal, n=5) and *LmexCen*^*−/−*^ *-*immunized (Imm chal, n=6) hamsters were determined after 10-12 months post *L. donovani* infected sand fly challenge. Results represent (the geometric means with 95% Cl) one experiment (*p- 0.01 and **p-0.008 values were determined by Mann-Whitney two-tailed test). Results are from one experiment.

### Immunization with *LmexCen*^*-/-*^ confers long-term protection against fatal visceral infection in a hamster model

Next, to determine the durability of protection initiated by *LmexCen*^*-/-*^ immunization, hamsters were challenged with virulent wild-type *L. donovani* through needle injection 8-months post-immunization (Fig. 5A). Age-matched naïve *L. donovani* infected animals were used as a control group (non-immune challenged group). Analysis of spleen and liver parasite loads after 8-months post-challenge showed significant control of spleen and liver parasite load. Immunized hamsters showed a ∼1.5 and a ∼1.0log-fold reduction in the spleen (Fig. 5B) and liver (Fig. 5C) parasite burden, respectively, compared to non-immunized challenged hamsters. These data confirm the long-lasting protective immunity of *LmexCen*^*-/-*^ parasites against visceral infection in the preclinical hamster model.

**Figure 5.**
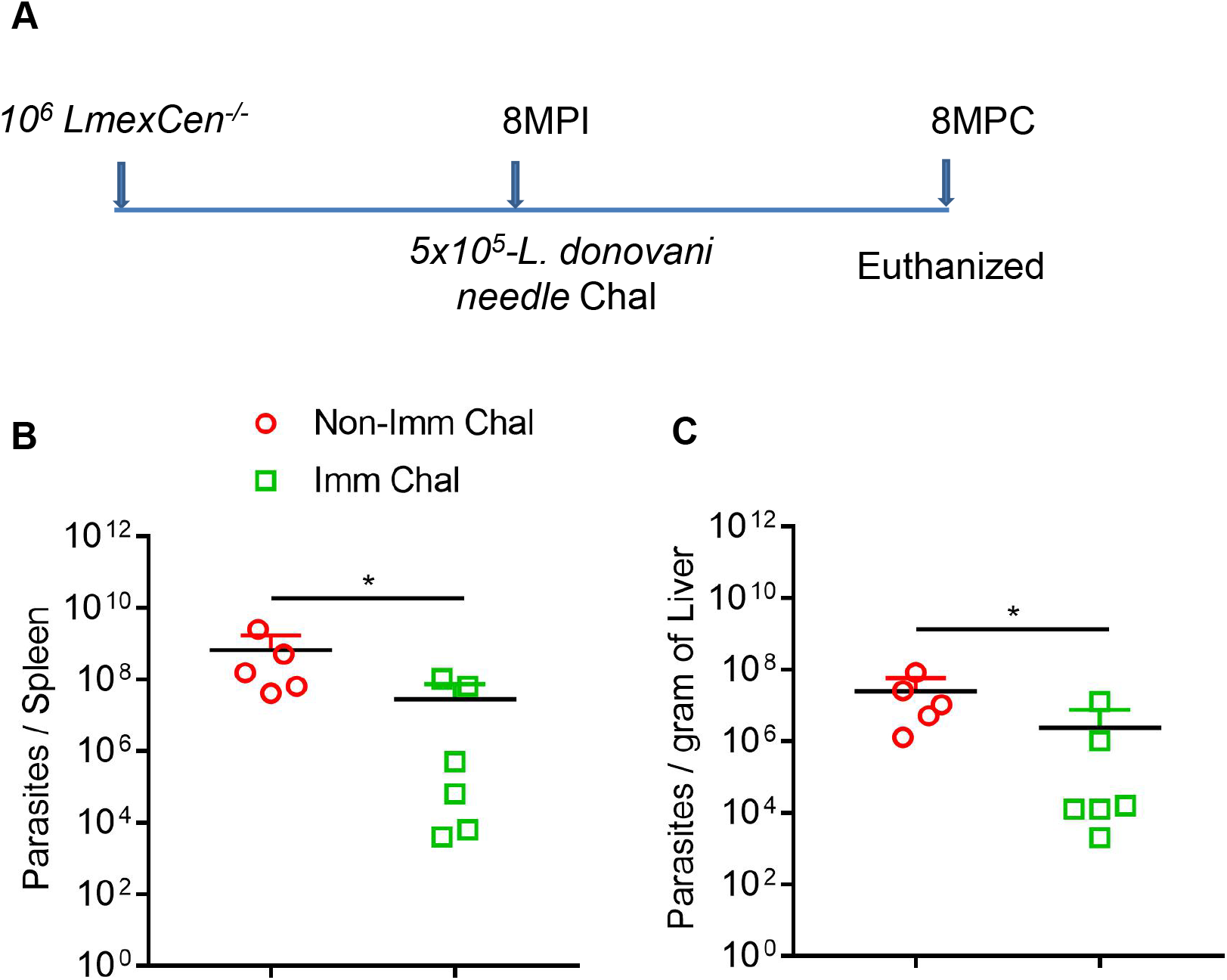
*LmexCen*^*−/−*^ immunization confers long term protection against needle challenge *L. donovani* infection in hamsters. (**A**) Schematic representation of experimental plan to determine the long-term protective efficacy of *LmexCen*^*-/-*^ parasites against *L. donovani* infection through needle injection. (**B** and **C**) Spleen (B) and Liver (C) parasite burden of *LmexCen*^*−/−*^ immunized (Imm Chal, n=6) and age matched non-immunized (Non-Imm Chal, n=5) hamsters were determined at 8 months post needle challenge. Results (means ± SD) are from one experiment (*p-0.03 values were determined by Mann Whitney two-tailed test). MPI, Months post-immunization; MPC, months post-challenge.

## DISCUSSION

Antileishmanial treatment is complex and shows toxic side effects. Therefore, a vaccine is urgently needed against human leishmaniasis, VL in particular. Epidemiological evidence and laboratory studies suggested that infection with one species of *Leishmania* species can cross-protect against other species ^8,14,15,28^. Particularly, recovery from infection with old world species such as *L. major* parasites has been shown to confer protection against New World species *L. infantum*, which causes fatal visceral leishmaniasis, confirming that leishmanization is a viable vaccination approach ^14^. Although, *Leishmania mexicana* induces skin pathology similar to *L. major* infection, infection with *L. mexicana* can result in the development of non-healing lesions in contrast to the spontaneously healing lesions observed in *L. major* infections ^18,19^. In addition, *L. mexicana* infection is associated with a dominant Th2 response as compared to *L. major* parasites ^20^, whereas the role of Th2 response in pathogenesis of *L. major* is debatable ^29,30^. Due to the differences in clinical severity and the immunological mechanisms of pathogenesis, a process analogous to leishmanization involving *L. major* infection in the old world as a means of acquiring protective immunity may not be feasible with new world species such as *L. mexicana*. Therefore, it may be important to develop a vaccine that can protect against infections of *Leishmania* species prevalent in the new world. Towards that goal, we have developed *centrin* gene-deleted *L. mexicana* parasites using the CRISPR/Cas9 technique. Our studies in mouse infection models indicated that *LmexCen*^*-/-*^ immunization confers protection against homologous wild-type *L. mexicana* challenge infection ^22^. Since both CL and VL are prevalent in Americas, a vaccine against these diseases will be needed. Therefore, we wanted to test the hypothesis whether a new world strain derived *LmexCen*^*-/-*^ can also protect against a VL infection.

This study reports that *LmexCen*^*-/-*^ parasites induce significant host immune response and protect hamsters from developing severe VL disease upon challenge with Old World *L. donovani* parasites. Our results show that *LmexCen*^*-/-*^ parasites do not cause any pathology in susceptible hosts such as hamsters ^31^. Our data further confirm the safety characteristics of *LmexCen*^*-/-*^ parasites in the hamster model, as was previously shown in mouse models of infection ^22^. Immunization with *LmexCen*^*-/-*^ showed downregulation of the Th2 response in hamster spleens compared to the spleens of *LmexWT*-infected animals, as was also observed in mouse models of infection ^22^. Specifically, while *LmexWT* inoculated hamsters developed a robust Th2 response, as evident by the IL-10 and IL-4 expression levels, *LmexCen*^***−/−***^ immunized hamsters displayed a diminished expression of both these cytokines relative to *LmexWT* infection. On the other hand, there was similar induction of pro-inflammatory cytokine IFN-γ, TNF−α and IL−12 between *LmexWT* infected and *LmexCen*^***−/−***^ immunized hamster spleens despite the significant difference in parasite burden in these groups. Overall, analysis of splenic immune response showed the ratios between IFN-γ/IL-10 and IFN-γ/ IL-4 higher in *LmexCen*^*-/-*^ immunized hamsters than in *LmexWT* infection. In mice infected with *L. major*, IL-4 can play a critical role in modulating the establishment of a protective Th1 response ^29,32^. However, in *L. mexicana* infection, IL-10 and IL-4 play an crucial role in disease susceptibility in mice models ^33,34^. In contrast, *LmCen*^*-/-*^ immunization induced a dominant Th1 response (60 folds high IFN-γ compared to *LmWT*) evident throughout the immunization period ^25^. However, IL-10 and IL-4 responses were significantly diminished in *LmexCen*^*-/-*^ immunization (6 folds enrichment of IFN-γ/IL-10 in *LmCen*^*-/-*^ versus 2-3 folds enrichment in *LmexCen*^*-/-*^), indicating that *LmCen*^*-/-*^ induces an exaggerated immune response when tested at an equivalent timepoint post-immunization in hamsters. It is also well established that in *L. mexicana* infection, a diminished expression of Th2 response is crucial to provide host protection ^35,36^. Our current studies suggest that immunization with *LmexCen*^***−/−***^ parasites in both rodent models induce a similar protective immune response.

Next, we observed that the immune response generated by *LmexCen*^***−/−***^ immunization induced significant host protection against *L. donovani* challenge both in the spleen and liver up to 15 months of post-challenge, as evident by the substantial reduction in parasite burden compared to non-immunized animals. In contrast, all non-immunized challenged hamsters developed severe symptoms of VL and succumbed to death by 15 months of post-challenge. On the contrary, 80 % of immunized and challenged animals survived until the study ended. In addition, *LmexCen*^***−/−***^ induced immunity was long-lasting, as evidenced by the parasite control in animals immunized over eight months compared to non-immunized animals. This could be due to the persistence of vaccine parasites in immunized animals that may play an essential role in maintaining a protective immune response. In support of that, we observed a small number of live parasites recovered from some of the hamsters even after 11 weeks post-immunization. However, it is unclear if, similar to *L. major* infection, the persistence of *L. mexicana* parasites is also necessary to maintain long-term protection, which requires further studies.

In conclusion, we have demonstrated that using CRISPR/Cas9 mediated centrin gene deletion of L. mexicana parasite, a new world Leishmania species, is safe and effective against CL and the severe form of VL caused by old world species, *L. donovani*. Furthermore, these studies demonstrate that centrin deleted parasites, whether from the old world or the new world, *Leishmania* offers an expanded number of vaccine candidates that could be developed to control leishmaniasis in all endemic regions of the world and eventually eliminate the disease.

## MATERIALS AND METHODS

### Study design and Ethical statement

Immunization and challenge infections were performed in a hamster model of VL to determine the efficacy of *centrin* gene-deleted *L. mexicana* (*LmexCen*^*-/-*^) parasites as a vaccine against experimental *L. donovani* infection. All animal experiments in this study were reviewed and approved by the Animal Care and Use Committee of the Center for Biologics Evaluation and Research, U.S. Food and Drug Administration (ASP 1999#23) and the National Institute of Allergy and Infectious Diseases (NIAID) (http://grants.nih.gov/grants/olaw/references/phspolicylabanimals.pdf) under animal protocol LMVR4E. The NIAID DIR Animal Care and Use Program complies with the Guide for the Care and Use of Laboratory Animals and with the NIH Office of Animal Care and Use and Animal Research Advisory Committee guidelines. The housing condition of animals were followed standard guidelines by NIH guidelines for the humane care and use of animals.

### Animals and parasites

Six to eight-week-old female outbred Syrian golden hamsters (*Mesocricetus auratus*) were obtained from the Harlan Laboratories Indianapolis, USA. All animals were housed either at the Food and Drug Administration (FDA) animal facility, Silver Spring (MD) or the National Institute of Allergy and Infectious Diseases (NIAID), Twin-brook campus animal facility, Rockville (MD), under pathogen-free conditions. The wild-type *L. donovani* (*LdWT*) (MHOM/SD/62/1S) parasites, wild-type *Leishmania mexicana* (MNYC/B2/62/m379) parasites (*LmexWT*) and *centrin* gene-deleted *LmexCen*^*-/-*^ promastigotes were cultured as previously described ^22,25^

### Immunization of hamsters and determination of parasite load

Six to eight-week-old female hamsters were immunized with 10^6^ total stationary-phase *LmexCen*^*-/-*^ parasites by intradermal injection in the left ear in 10μl PBS using a 29-gauge needle (BD Ultra-Fine). In addition, the age-matched control group of animals were infected with 10^6^ total stationary-phase *L. mexicana* wild-type (*LmexWT*) promastigotes. Lesion size was monitored weekly by measuring the diameter of the ear lesion using a direct reading vernier calliper. Parasite burdens in the ear, draining lymph node (dLN), spleen, liver and bone marrow tissues were determined by limiting dilution assay as described in previous studies ^25^.

### Determination of cytokine expression in splenocytes by real-time PCR

Single cell suspensions from the spleens were made in hamsters at eleven weeks post-immunization with *LmexCen*^*-/-*^ as well as infected with *LmexWT*. Then they were stimulated with *L. mexicana* freeze-thawed antigen (FTAg), and total RNA was extracted using PureLink RNA Mini kit (Ambion) at 16h after restimulation. Aliquots (400ng) of total RNA were reverse transcribed into cDNA by a high-capacity cDNA reverse transcription kit using random hexamers (Applied Biosystems). Cytokine gene expression levels were determined by TaqMan probe (TaqMan, Universal PCR Master Mix, Applied Biosystems) PCR using a CFX96 Touch Real-Time System (BioRad, Hercules, CA). The sequences of the primers (forward and reverse) and probes (5′ 6-FAM and 3′ TAMRA Quencher) were used to detect the gene expression described as before ^25^. The data were analyzed with CFX Manager Software. The expression levels of genes of interest were determined by the 2^−ΔΔCt^ method; samples were normalized to γ-actin expression and determined relative to expression values from naive hamsters.

### *LmexCen*^*-/-*^ Immunization and challenge with needle infection

Six to eight-week-old female hamsters were immunized with 10^6^ total stationary-phase *LmexCen*^*-/-*^parasites by intradermal injection in the left ear in 10μl PBS using a 29-gauge needle (BD Ultra-Fine). Eleven weeks post-immunization, animals were challenged with a needle injection (ID route) with virulent 5 × 10^5^ metacyclic *L. donovani* (Ld1S) promastigotes into the contralateral ear. Age-matched non-immunized hamsters were similarly challenged and used as a control group. Hamsters were monitored daily, and after various periods of post-challenge (9- and 15-month post-challenge), animals were sacrificed, and the parasite load in organs (spleen and liver) was measured by limiting dilution as previously described ^25^.

### *LmexCen*^*-/-*^ Immunization and challenge with sand fly mediated infection

Female 2- to 4-day-old *Lutzomyia longipalpis* sand flies were infected with *L. donovani* amastigotes (freshly isolated from a sick hamster) as described previously ^25^. Parasite loads and percentage of metacyclic per midgut were determined using hemocytometer counts. Thirty sand flies on day 15- after infection were applied to the ear of each *LmexCen*^***−/−***^ immunized and age-matched non-immunized hamsters used for subsequent transmission as described previously. The number of blood-fed flies was determined after transmission as a qualitative measurement. Each hamster received an average of 18-infected bites per transmission. Due to COVID related restrictions it was not possible to have separate control groups for *LmCen*^-/-^ and *LmexCen*^-/-^ studies that were conducted concurrently. Thus, the sand fly challenge experiments with non-immunized hamster controls shown in the current study were performed alongside our previous studies on *LmCen*^*-/-*^ parasites ^25^. Hamsters were monitored daily during infection, and after 9-15months of post-challenge, animals were sacrificed, and parasite load in organs (spleen and liver) was measured by limiting dilution as previously described ^25^.

### Statistical analysis

Statistical analysis of differences between groups was determined by unpaired two-tailed Mann–Whitney t-test using Graph Pad Prism 7.0 software.

### Data availability statement

The data that support the findings of this study are available from the corresponding author upon reasonable request.

## Acknowledgments

Funding was provided from the Global Health Innovative Technology Fund, the Canadian Institutes of Health Research (to GM), intramural funding from CBER, FDA (to HLN), and the Fonds de recherche du Québec – Santé (to PL). This research was supported, in part, by the Intramural Research Program of the NIH, National Institute of Allergy and Infectious Diseases (F.O., J.O., C.M, S.K. and J.G.V.). The findings of this study are an informal communication and represent the authors’ own best judgments. These comments do not bind or obligate the Food and Drug Administration.

## Author contributions

SK (Subir Karmakar), GV, WWZ, PL, NI, FO, JO, SG, CM and RD conducted experiments, analyzed data and helped to write the manuscript. SK (Subir Karmakar), RD, ARS, SH, PD, SK (Shaden Kamhawi), SS, JGV and HLN designed experiments, analyzed data and wrote the manuscript.

## REFERENCES

1 Burza, S., Croft, S. L. & Boelaert, M. Leishmaniasis. Lancet 392, 951–970, doi:10.1016/S0140-6736(18)31204-2 (2018).

2 Kaye, P. & Scott, P. Leishmaniasis: complexity at the host-pathogen interface. Nat Rev Microbiol 9, 604–615, doi:10.1038/nrmicro2608 (2011).

3 McGwire, B. S. & Satoskar, A. R. Leishmaniasis: clinical syndromes and treatment. QJM 107, 7–14, doi:10.1093/qjmed/hct116 (2014).

4 Taslimi, Y., Zahedifard, F. & Rafati, S. Leishmaniasis and various immunotherapeutic approaches. Parasitology 145, 497–507, doi:10.1017/S003118201600216X (2018).

5 Ghorbani, M. & Farhoudi, R. Leishmaniasis in humans: drug or vaccine therapy? Drug Des Devel Ther 12, 25–40, doi:10.2147/DDDT.S146521 (2018).

6 Ponte-Sucre, A. et al. Drug resistance and treatment failure in leishmaniasis: A 21st century challenge. PLoS Negl Trop Dis 11, e0006052, doi:10.1371/journal.pntd.0006052 (2017).

7 Lainson, R. & Shaw, J. J. Leishmaniasis in Brazil: XII. Observations on cross-immunity in monkeys and man infected with Leishmania mexicana mexicana, L. m. amazonensis, L. braziliensis braziliensis, L. b. guyanensis and L. b. panamensis. J Trop Med Hyg 80, 29–35 (1977).

8 Porrozzi, R., Teva, A., Amaral, V. F., Santos da Costa, M. V. & Grimaldi, G., Jr. Cross-immunity experiments between different species or strains of Leishmania in rhesus macaques (Macaca mulatta). Am J Trop Med Hyg 71, 297–305 (2004).

9 Ostyn, B. et al. Incidence of symptomatic and asymptomatic Leishmania donovani infections in high-endemic foci in India and Nepal: a prospective study. PLoS Negl Trop Dis 5, e1284, doi:10.1371/journal.pntd.0001284 (2011).

10 Jeronimo, S. M. et al. Natural history of Leishmania (Leishmania) chagasi infection in Northeastern Brazil: long-term follow-up. Clin Infect Dis 30, 608–609, doi:10.1086/313697 (2000).

11 Nadim, A., Javadian, E., Tahvildar-Bidruni, G. & Ghorbani, M. Effectiveness of leishmanization in the control of cutaneous leishmaniasis. Bull Soc Pathol Exot Filiales 76, 377–383 (1983).

12 Kellina, O. I. Problem and current lines in investigations on the epidemiology of leishmaniasis and its control in the U.S.S.R. Bull Soc Pathol Exot Filiales 74, 306–318 (1981).

13 Seyed, N., Peters, N. C. & Rafati, S. Translating Observations From Leishmanization Into Non-Living Vaccines: The Potential of Dendritic Cell-Based Vaccination Strategies Against Leishmania. Front Immunol 9, 1227, doi:10.3389/fimmu.2018.01227 (2018).

14 Romano, A., Doria, N. A., Mendez, J., Sacks, D. L. & Peters, N. C. Cutaneous Infection with Leishmania major Mediates Heterologous Protection against Visceral Infection with Leishmania infantum. J Immunol 195, 3816–3827, doi:10.4049/jimmunol.1500752 (2015).

15 Zijlstra, E. E., el-Hassan, A. M., Ismael, A. & Ghalib, H. W. Endemic kala-azar in eastern Sudan: a longitudinal study on the incidence of clinical and subclinical infection and post-kala-azar dermal leishmaniasis. Am J Trop Med Hyg 51, 826–836 (1994).

16 Okwor, I. & Uzonna, J. Vaccines and vaccination strategies against human cutaneous leishmaniasis. Hum Vaccin 5, 291–301, doi:10.4161/hv.5.5.7607 (2009).

17 Noazin, S. et al. First generation leishmaniasis vaccines: a review of field efficacy trials. Vaccine 26, 6759–6767, doi:10.1016/j.vaccine.2008.09.085 (2008).

18 Scorza, B. M., Carvalho, E. M. & Wilson, M. E. Cutaneous Manifestations of Human and Murine Leishmaniasis. Int J Mol Sci 18, doi:10.3390/ijms18061296 (2017).

19 Gabriel, A. et al. Cutaneous Leishmaniasis: The Complexity of Host’s Effective Immune Response against a Polymorphic Parasitic Disease. J Immunol Res 2019, 2603730, doi:10.1155/2019/2603730 (2019).

20 Rosas, L. E. et al. Genetic background influences immune responses and disease outcome of cutaneous L. mexicana infection in mice. Int Immunol 17, 1347–1357, doi:10.1093/intimm/dxh313 (2005).

21 Paiva, M. B. et al. A Cytokine Network Balance Influences the Fate of Leishmania (Viannia) braziliensis Infection in a Cutaneous Leishmaniasis Hamster Model. Front Immunol 12, 656919, doi:10.3389/fimmu.2021.656919 (2021).

22 Volpedo, G. et al. Centrin-deficient Leishmania mexicana confers protection against New World cutaneous leishmaniasis. NPJ Vaccines 7, 32, doi:10.1038/s41541-022-00449-1 (2022).

23 Selvapandiyan, A. et al. Centrin gene disruption impairs stage-specific basal body duplication and cell cycle progression in Leishmania. J Biol Chem 279, 25703–25710, doi:10.1074/jbc.M402794200 (2004).

24 Selvapandiyan, A. et al. Centrin1 is required for organelle segregation and cytokinesis in Trypanosoma brucei. Mol Biol Cell 18, 3290–3301, doi:10.1091/mbc.e07-01-0022 (2007).

25 Karmakar, S. et al. Preclinical validation of a live attenuated dermotropic Leishmania vaccine against vector transmitted fatal visceral leishmaniasis. Commun Biol 4, 929, doi:10.1038/s42003-021-02446-x (2021).

26 Zhang, W. W. et al. A second generation leishmanization vaccine with a markerless attenuated Leishmania major strain using CRISPR gene editing. Nat Commun 11, 3461, doi:10.1038/s41467-020-17154-z (2020).

27 Aslan, H. et al. A new model of progressive visceral leishmaniasis in hamsters by natural transmission via bites of vector sand flies. J Infect Dis 207, 1328–1338, doi:10.1093/infdis/jis932 (2013).

28 Dey, R. et al. Characterization of cross-protection by genetically modified live-attenuated Leishmania donovani parasites against Leishmania mexicana. J Immunol 193, 3513–3527, doi:10.4049/jimmunol.1303145 (2014).

29 Biedermann, T. et al. IL-4 instructs TH1 responses and resistance to Leishmania major in susceptible BALB/c mice. Nat Immunol 2, 1054–1060, doi:10.1038/ni725 (2001).

30 Hurdayal, R. et al. Deletion of IL-4 receptor alpha on dendritic cells renders BALB/c mice hypersusceptible to Leishmania major infection. PLoS Pathog 9, e1003699, doi:10.1371/journal.ppat.1003699 (2013).

31 Wilson, H. R., Dieckmann, B. S. & Childs, G. E. Leishmania braziliensis and Leishmania mexicana: experimental cutaneous infections in golden hamsters. Exp Parasitol 47, 270–283, doi:10.1016/0014-4894(79)90079-1 (1979).

32 Poudel, B. et al. Acute IL-4 Governs Pathogenic T Cell Responses during Leishmania major Infection. Immunohorizons 4, 546–560, doi:10.4049/immunohorizons.2000076 (2020).

33 Padigel, U. M., Alexander, J. & Farrell, J. P. The role of interleukin-10 in susceptibility of BALB/c mice to infection with Leishmania mexicana and Leishmania amazonensis. J Immunol 171, 3705–3710, doi:10.4049/jimmunol.171.7.3705 (2003).

34 Alexander, J. et al. An essential role for IL-13 in maintaining a non-healing response following Leishmania mexicana infection. Eur J Immunol 32, 2923–2933, doi:10.1002/1521-4141(2002010)32:10<2923::AID-IMMU2923>3.0.CO;2-E (2002).

35 Bryson, K. J. et al. BALB/c mice deficient in CD4 T cell IL-4Ralpha expression control Leishmania mexicana Load although female but not male mice develop a healer phenotype. PLoS Negl Trop Dis 5, e930, doi:10.1371/journal.pntd.0000930 (2011).

36 Satoskar, A., Bluethmann, H. & Alexander, J. Disruption of the murine interleukin-4 gene inhibits disease progression during Leishmania mexicana infection but does not increase control of Leishmania donovani infection. Infect Immun 63, 4894–4899, doi:10.1128/iai.63.12.4894-4899.1995 (1995).

